## Supplementary Materials for "DeepRank: A deep learning framework for data mining 3D protein-protein interfaces"

### DeepRank:

**1 Netherlands eScience Center, Science Park 140, 1098 XG, Amsterdam, The Netherlands**

**2 Bijvoet Centre for Biomolecular Research, Faculty of Science - Chemistry, Utrecht University, Padualaan 8, 3584 CH Utrecht, The Netherlands**

**3 Center for Molecular and Biomolecular Informatics, Radboudumc, Greet Grooteplein 26-28, 6525 GA Nijmegen, The Netherlands**

#### 1 - Software Architecture

DeepRank is built as a comprehensive Python3 package that allows end-to-end classification and/or ranking of protein-protein interfaces. The package is accessible on GitHub: https://github.com/DeepRank/deeprank. DeepRank and its dependencies can easily be installed using the PyPI package manager using the command:

```
pip install deeprank
```

The global architecture of the software is represented in **Fig. S1**. The two main components of DeepRank are respectively in charge of the feature calculation and of the training of the models. These two components communicate via dedicated HDF5 files containing the mapped features that are used as input for the training. Detailed descriptions on how to use the code can be found in the online documentation: https://deeprank.readthedocs.io/

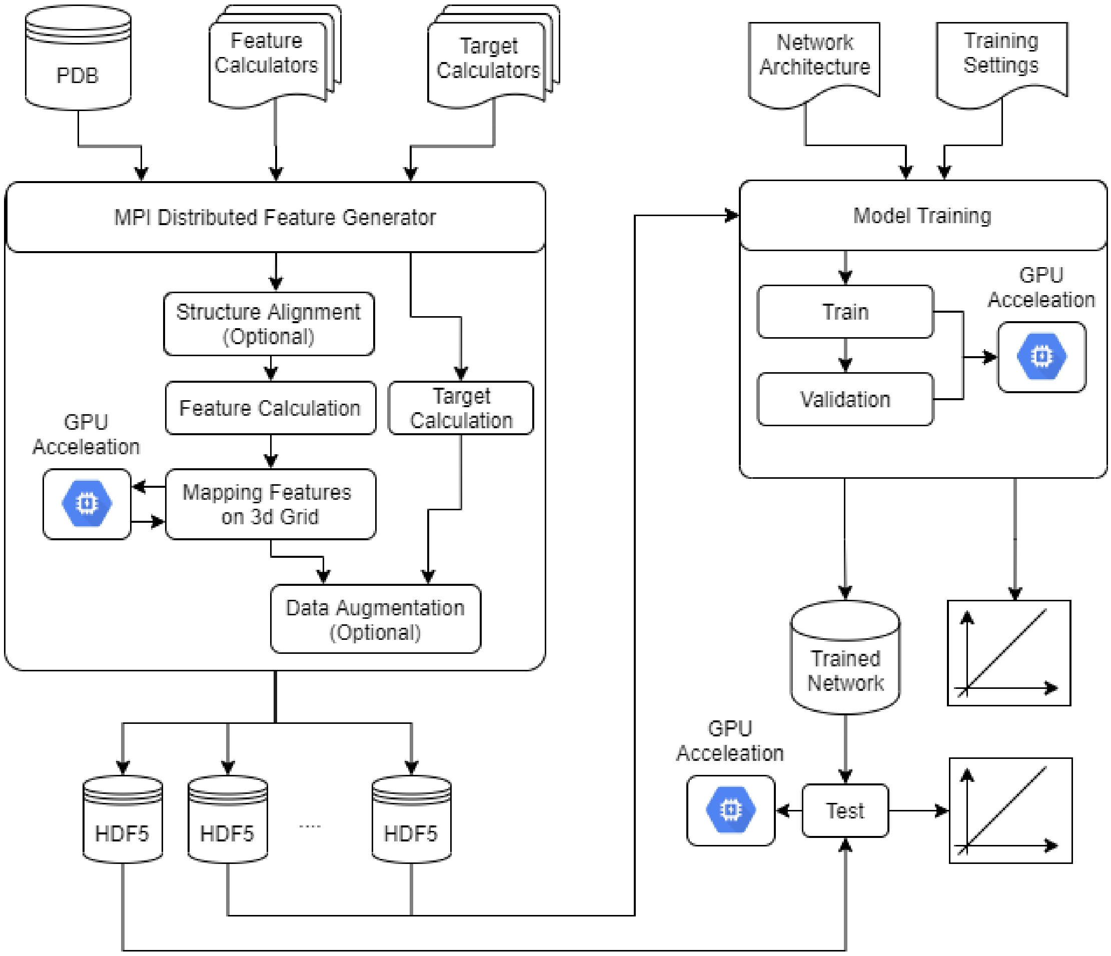

**Fig S1. Architecture of the DeepRank software.** The two main blocks are respectively in charge of 1) 3D feature grid generation and 2) training and evaluating of the network. The package contains two main parts: one dedicated to the generation of 3D feature grids and the other dedicated to the training of neural networks.

#### 1-A Data Generation Module

The data generating module takes as inputs the PDB files of the complexes/models used for training/validation/testing. The features listed in **Table 1** and targets (e.g., class labels in the context of classification, or binding affinities in the context of affinity predictions) presented in the main text are all already implemented in DeepRank and can simply be called while instantiating the DataGenerator object. If users need new features and/or targets, these can be easily added to be used by the data generator. A tutorial explaining this process is available in the online documentation:

https://deeprank.readthedocs.io/en/latest/advTuto.html#

Once PDB files, features and targets metrics are specified, the data generator will split the processing of the PDB files among the requested number of MPI processes. This can greatly accelerate the data pre-processing and featurization process. Since it is essentially embarrassingly parallel it scales linearly with the number of MPI processes. For each PDB file, the feature values will be first calculated. These values are localized on a given atom or residue. They are subsequently mapped on a 3D Grid using a Gaussian mapping (see Methods in the main manuscript).

The mapping process is supported by MPI distributed processes and GPU offloading through dedicated CUDA kernels implemented in DeepRank to ensure efficient computations for very large datasets. As an indication of the processing time, mapping one feature on a 30x30x30 grid requires 0.135 s on a single CPU (Intel(R) Xeon(R) CPU E5-2650 v4 @ 2.20GHz) compared to 0.085 s on an NVIDIA GeForce Ti1080. A larger speed-up is expected for grids containing more points. Note that, in order to reduce the size of the HDF5 files, the 3D grids can also be computed on demand during the training of the model, for example to change the grid size and/or resolution. This however comes, at the price of increased computational costs for training a model. The target values are also calculated using either the native methods implemented in DeepRank or user-defined ones.

Data augmentation is supported in DeepRank by randomly rotating the 3D structure of the complex around the geometric center of its interface, after which features are automatically mapped onto the grid. For situations where random orientations of the PPIs are not desired, protein complexes can be aligned by DeepRank using Principal Component Analysis (PCA)-based alignments along cartesian axes. Users can specify a given number of copies for each structure: Each copy will be randomly rotated to augment the data set and expose the network to different orientations of the same interface.

All resulting feature and target data are stored in a series of HDF5 files. The data set is split into multiple HDF5 files, one file for each MPI process used. Hence if one compute the features of docking models of 1 given complex using 24 MPI process, 24 HDF5 files will be created each containing the features of a subset of docking conformations. Using HDF5 files not only reduces the memory requirements for the mapped features through the native compression schemes of HDF5 but also allows to easily explore the data sets using our in-house HDF5 browser DeepXplorer specifically tuned for the data generated here (https://github.com/DeepRank/DeepXplorer). This graphical user interface allows to easily browse and visualize the mapped features through popular molecular viewers such as VMD and PyMol.

##### 1-B Training Module

The module in charge of the training of the architecture can easily read the data contained in the HDF5 files. They can specify which HDF5 file to use in training/validation and testing and can also select complexes satisfying defined criteria such as, for example, models with iRMSD larger and/or lower than a given threshold. The training module also needs as input a Python file containing the architecture of the neural network to be used. An example file is provided with the package (i.e., `deeprank/learn/model3d.py`, which contains simple 3D CNN architectures for classification and regression) but users can easily expand on this architecture to optimize the hyper-parameters of the model.

The data set resulting from the selection of a subset of conformations is then used to train the model. Note that the dataset is not loaded in memory and instead each data point is dynamically loaded when needed in the minibatch. At the end of the training, a plot of losses over epoches is generated assisting the user to identify the optimal network parameters. The best models with the lowest losses are output together with the training data in a dedicated HDF5 file. This allows easy exploration of the training and validation performance through the HDF5 browser. This plot can then be used on an independent part of the total data set for testing purposes. For classification, DeepRank outputs boxplots of prediction scores and accuracy plots for train/validation/test. For regression, DeepRank outputs scatter plots for predicted values vs. target values.

#### 2 - Classification of biological interfaces vs. crystal interfaces

We provide here details about the architecture and training procedure used for the classification of biological interfaces vs. crystal ones.

##### 2-A Network architecture

The network used for this task is summarized in **Fig. S2**. The network contains two 3D CNN/Max pooling blocks followed by two linear fully connected layers. This results in 117,362 learnable parameters. 80% of 5739 complexes from the MANY data set were used as training set. Each complex was augmented with 30 rotations leading to a total of 142352 conformations. The remaining 20% of the MANY data set, i.e.1147 conformations were used in the validation data set. The test set was composed of the 161 conformations from an independent data set, namely the DC dataset. PSSM were used as features for residues in each chain (i.e., each residue is represented as 20 by 1 vector), leading to a total of 40 input channels.

```
Data Set Info:
Augmentation          : 30 rotations
Training set          : 142352 conformations
Validation set        : 1147 conformations
Test set              : 161 conformations
Number of channels    : 40
Grid Size             : 10, 10, 10
----------------------------------------------------------------
        Layer (type)               Output Shape         Param #
================================================================
            Conv3d-1          [-1, 80, 9, 9, 9]          25,680
         MaxPool3d-2          [-1, 80, 4, 4, 4]               0
            Conv3d-3         [-1, 120, 3, 3, 3]          76,920
         MaxPool3d-4         [-1, 120, 1, 1, 1]               0
            Linear-5                  [-1, 120]          14,520
            Linear-6                    [-1, 2]             242
================================================================
Total params: 117,362
Trainable params: 117,362
Non-trainable params: 0
```

**Fig S2. Architecture of the neural network and dataset size used for classification of biological vs. crystal interfaces.**

#### 2-B Loss and early stopping

**Fig. S3** shows the training and validation losses over epochs. As the validation loss is lowest at epoch 1, we used the trained network at epoch 1 as our final network. Note that epoch index starts with 0 thus epoch 1 means the network was trained by two rounds of the data. In addition, with a minibatch size of 8, and a total of 142352 conformations in the training set, 35588 individual optimization steps were performed at the end of the second epoch. We then applied the trained network on an independent set, the DC set, achieving 86% accuracy (see **Fig. 2**).

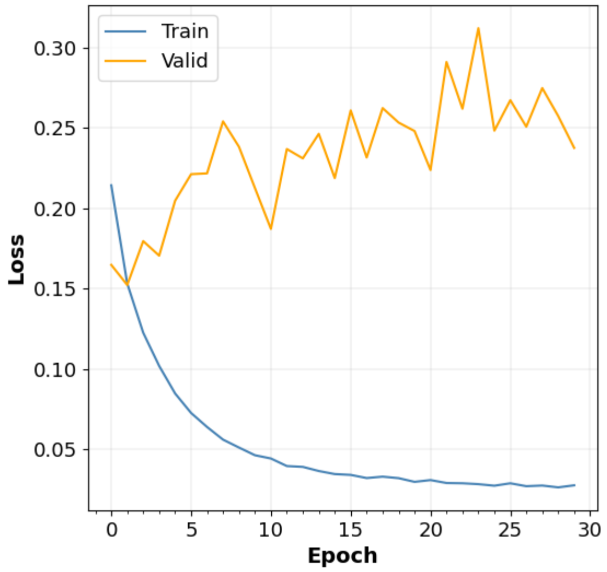

**Fig S3. Losses of training and validation on the MANY dataset for classification of biological vs crystal interfaces.**

#### 3 - Ranking docking models

In this section we present the details about the methods we used to train the neural network for the ranking problem as well as additional results.

##### 3-A Data selection

As mentioned earlier, the MPI-supported feature grid generation produces a series of HDF5 files. We ran the data generation using 24 MPI processes leading to 24 HDF5 files for each complex. For example, the 25000 docking conformations of PDB 3AAD were split into 24 HDF5 files, named 000_3AAD, 001_3AAD, …. 024_3AAD, each file containing all the mapped features, targets, etc  of about 1000 different docking conformations. As the conformations were not clustered, a large degree of redundancy exists between the conformations stored in the different files. After experimentation we concluded that considering a subset of models consisting of 3 HDF5 files per complex was sufficient to capture the distribution of the whole dataset (001, 002 and 003, ~0.4 million models). **Fig. S4** shows the distribution of iRMSD values of the whole dataset (gray shaded area) and of the portion used during our experiments. The left panel shows the distribution aggregated for all complexes while the right panel shows individual cases.

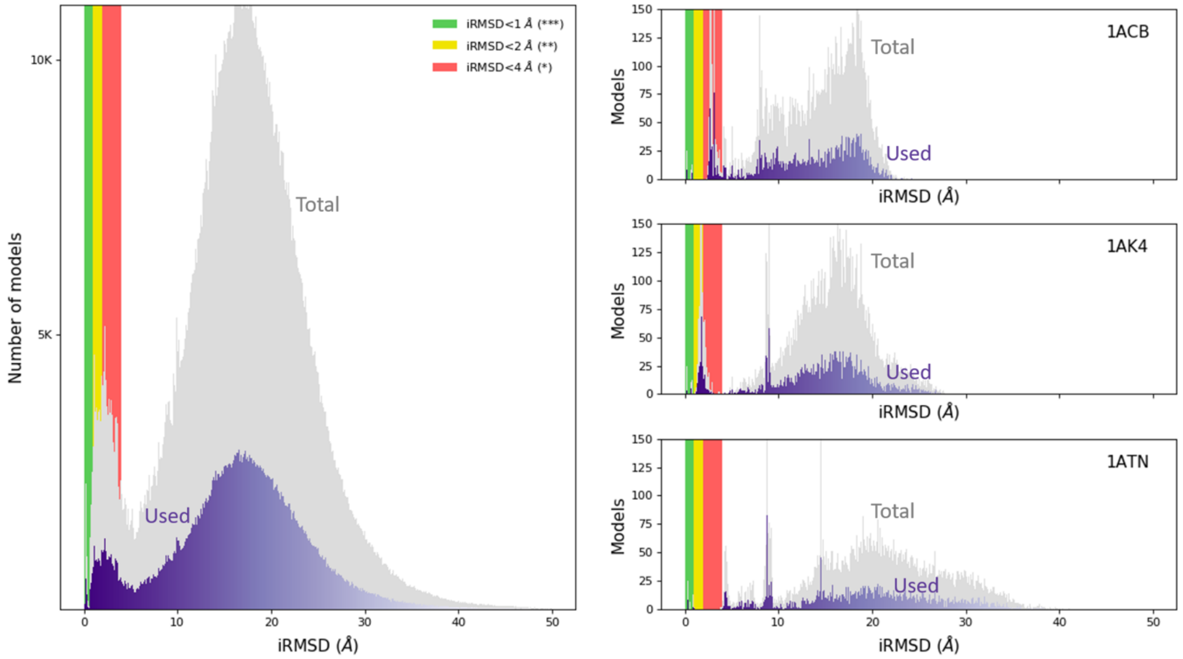

**Fig S4. Distribution of the iRMSD values.** The gray shaded area shows the distribution for the whole data set, while the purple area displays the distribution of the limited portion used during our experiments. The selected data reliably represents not only the entire dataset (left panel) but also individual cases (right panel). The histograms of the total dataset have been scaled by a factor ½ for clarity.

#### 3-B Neural network architecture

The network used for the ranking problem is summarized in **Fig. S5**. The network was composed for 8 sequential layers: a succession of 3D CNN, max-pooling and 3D batch normalization with 2 fully connected layers. It contains 7232 trainable parameters with a size of 6 MB.

```
Data Set Info:
Training set              : 338389 conformations
Augmentation              : 0 rotations
Validation set            : 40410 conformations
Test set                  : 39425 conformations
Number of channels        : 36
Grid Size                 : 30, 30, 30
----------------------------------------------------------------
        Layer (type)               Output Shape         Param #
================================================================
       BatchNorm3d-1      [-1, 36, 30, 30, 30]              72
            Conv3d-2       [-1, 6, 28, 28, 28]           5,838
       BatchNorm3d-3       [-1, 6, 28, 28, 28]              12
         MaxPool3d-4          [-1, 6, 9, 9, 9]               0
            Conv3d-5          [-1, 6, 7, 7, 7]             978
       BatchNorm3d-6          [-1, 6, 7, 7, 7]              12
         MaxPool3d-7          [-1, 6, 2, 2, 2]               0
            Linear-8                   [-1, 6]             294
       BatchNorm1d-9                   [-1, 6]              12
           Linear-10                   [-1, 2]              14
================================================================
Total params: 7,232
Trainable params: 7,232
Non-trainable params: 0
```

**Fig. S5: Architecture of the neural network and dataset size for the docking scoring problem.**

The performances reported in **Fig. 3** (and **Fig. S6**) were achieved in one epoch on 10-fold cross validation. For each epoch the training took 2 hours on a 16-core CPU and 1 GPU cards. We used ~340K models for training, 40K for validation and 40K for testing. We used the full set of 36 physico-chemical features (channels) that are predefined in DeepRank (**Table 1**).

#### 3-C Performance comparison between HADDOCK and DeepRank on BM5

**Fig. S6** shows the Hit Rate obtained with DeepRank and HADDOCK Score on models generated in the rigid-body docking stage only using the HADDOCK-it0 scoring function (left panel) and HADDOCK-itw scoring function (right panel). Using HADDOCK-it0 scoring function on models generated in the rigid-body docking phase leads to similar performance than DeepRank. However, using HADDOCK-itw scoring function on the same models, i.e. rigid-body docking models, leads to significantly lower performance. This illustrates the robustness of DeepRank on different model qualities.

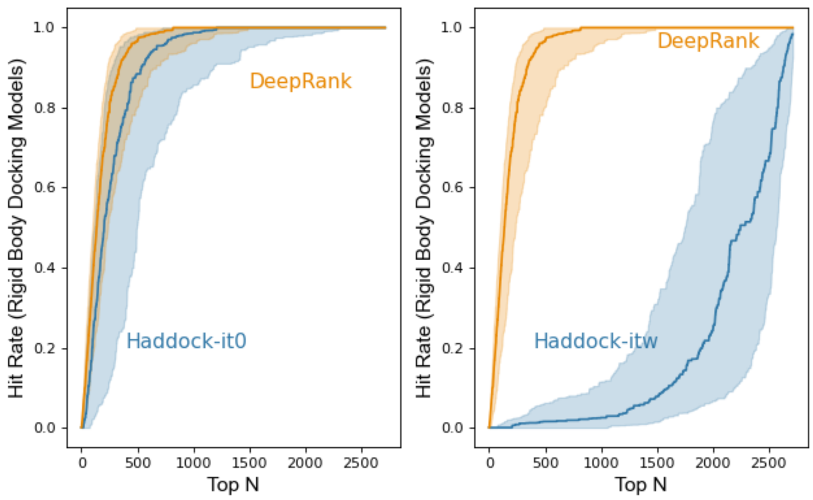

**Fig. S6. Comparing DeepRank and different HADDOCK scoring functions on rigid-body docking models.** DeepRank and HADDOCK-it0 show similar performance on these models (left). However HADDOCK-itw does not perform well on rigid-body-docking models (right).

We further checked the performance of DeepRank and HADDOCK Score on models originating from different docking stages: rigid-body docking, semi-flexible docking, and water refinement (**Fig. S7**). We used the most suitable HADDOCK scoring function in each case i.e. it0 for rigid-body docking models, it1 for semi-flexible docking models and itw for water refined models. DeepRank and HADDOCK show comparable performance with a slight advantage for DeepRank in each case.

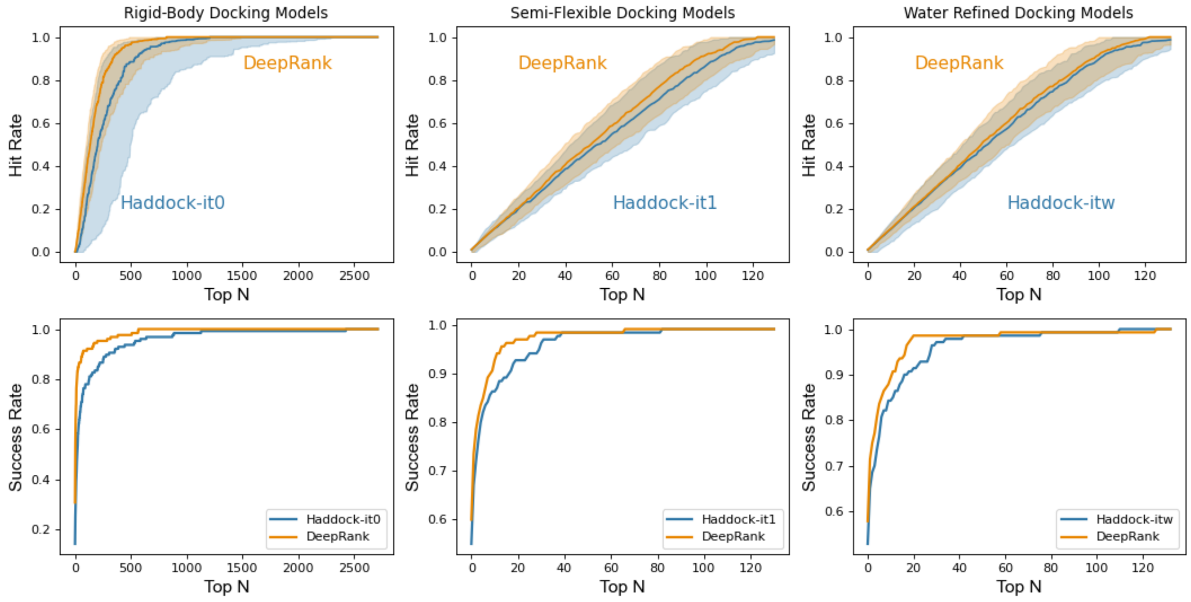

**Fig. S7. Ranking performance of DeepRank and HADDOCK.** Hit Rate over complex (top row) and Success Rate (bottom row), on models generated in different docking stages; rigid-body docking models (left); semi-flexible docking models (middle); water-refined models (right).

#### 3-D Performance comparison between HADDOCK and DeepRank on CAPRI

To further test the performance of DeepRank we have trained a final 3D CNN model using the conformations of all the 142 complexes and applied it to 13 cases from the CAPRI score set[1]. The CAPRI score set was generated by various docking software and represents an independent test set. We compared the DeepRank results to two leading scoring functions, the HADDOCK scoring function and the recently developed iScore[2,3], a graph-kernel based scoring function. To ensure optimal performance of the HADDOCK scoring function, the models of the CAPRI data set were subjected to a short energy minimization to remove clashes produced by rigid-body docking methods.

DeepRank is generally competitive with HADDOCK and iScore (**Fig. S8**), outperforming them on some cases especially in the top 200 and beyond (**Table S1**). DeepRank also performs very well when only a limited number of near-native models are present in the data set as is the case for T30 and T35 (**Fig. S8**) (2 out of 1343 for T30 and 3 out of 499 for T35, respectively). This suggests the ability of DeepRank to correctly identify favorable interactions that are ignored by the other methods and indicated a possible complementarity of these approaches.

The complementarity offered by DeepRank is further illustrated by comparing the rankings given by DeepRank and HADDOCK per cases as represented in **Fig. S9**. The lower a ranking the more likely HADDOCK/DeepRank considers a model to be near-native. As seen in these figures, the rankings given by DeepRank and HADDOCK can be significantly different. In cases of T40, T47, T50 and T53, a cluster of near-native conformations appears on the top left corner. This shows that DeepRank correctly ranks these conformations while HADDOCK mistakenly predicts them as wrong models. As an example, conformations of T50 are illustrated in **Fig. S10**. We can see that these conformations only differ by variations of the orientation of one chain with respect of the other one. This illustrates the large variability of the different scoring functions with respect to structural changes.

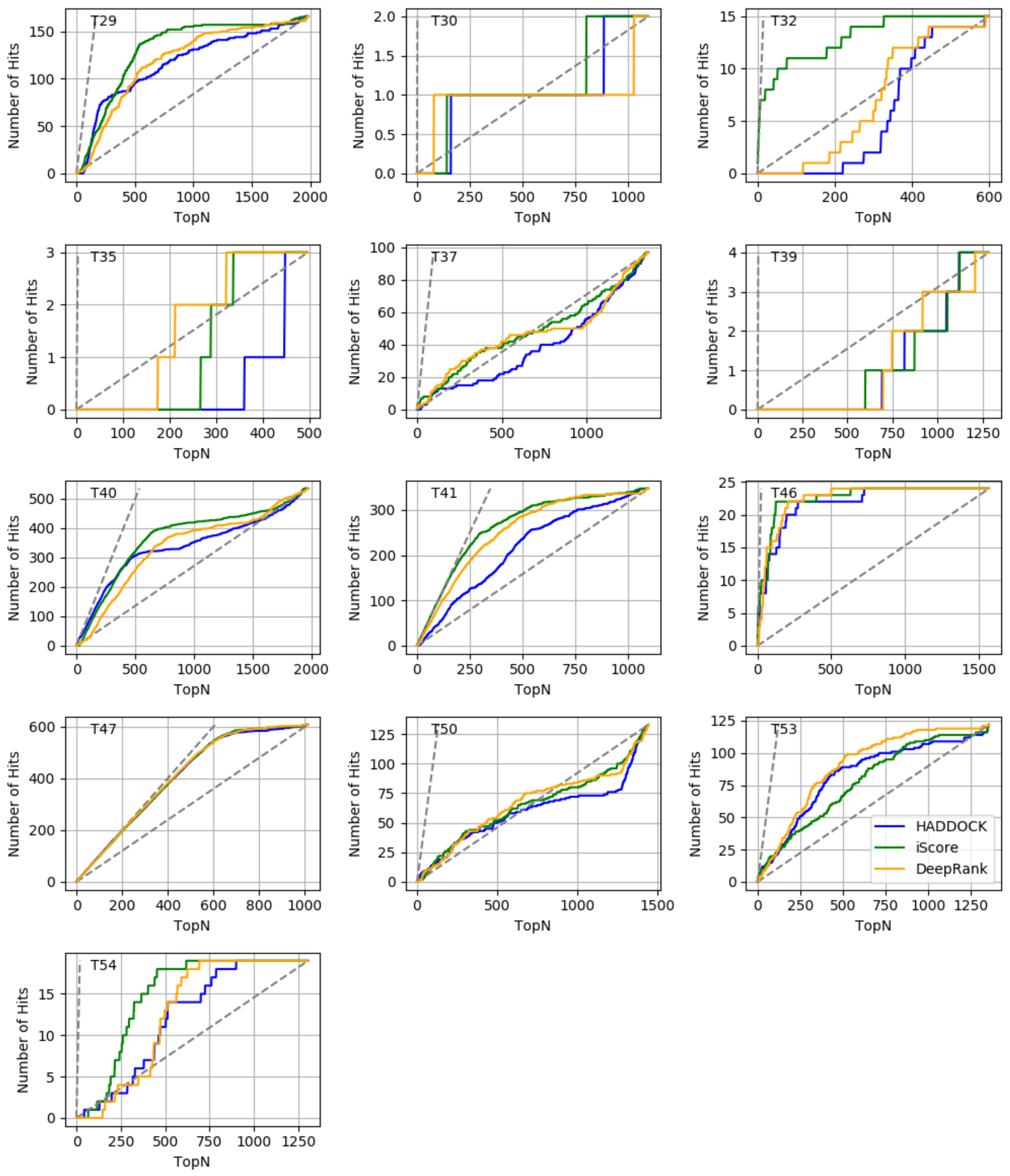

**Fig S8: Hit Rate plot of the 13 CAPRI cases given by HADDOCK, iScore and DeepRank.** DeepRank is competitive with these two scoring functions.

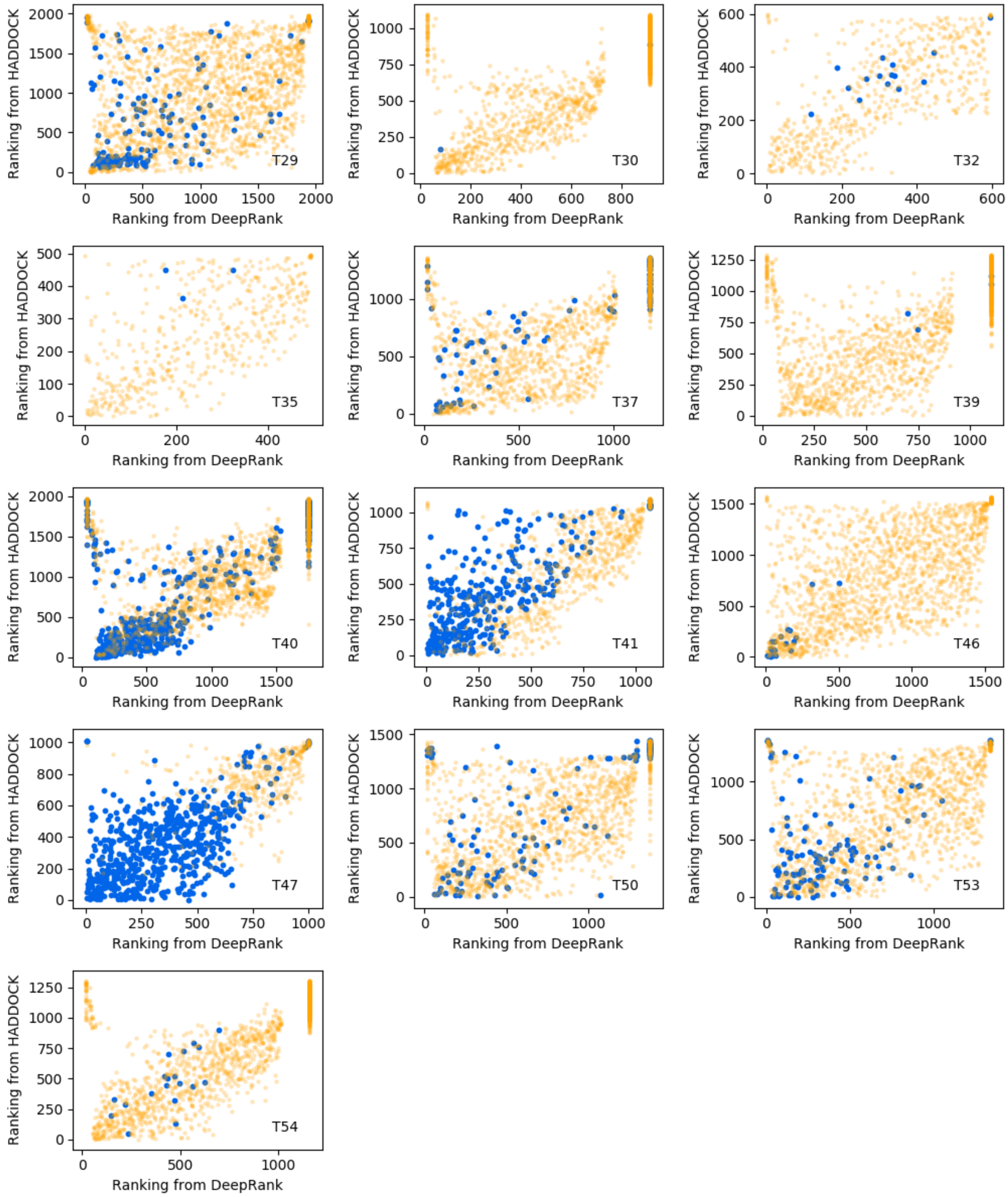

**Fig. S9: Comparison of the rankings obtained with HADDOCK and DeepRank on the CAPRI score set.** The lower a ranking the more likely HADDOCK/DeepRank considers a model to be near-native. Blue points represent near-native models while orange ones mark wrong models. The cluster of blue dots in the top-left corner is near-native models correctly identified by DeepRank but missed by HADDOCK. Inversely, the blue dots in the lower-right corner are near-native models that are correctly identified only by HADDOCK. Additionally, clusters of near-native models in the top right corner are those that are misclassified as wrong models by both DeepRank and HADDOCK.

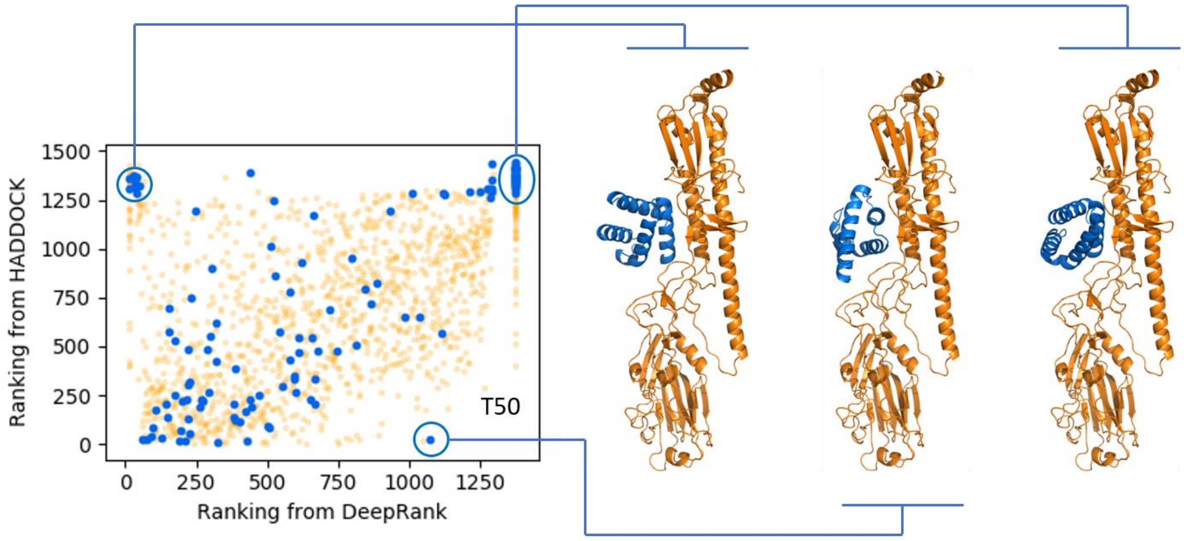

**Fig S10: Comparison of the rankings given by HADDOCK and DeepRank for the conformations of the T50 case.** The blue/orange dots mark near-native/wrong models. This scatter plot illustrates near-native conformations that are correctly or incorrectly ranked by either or both of the two scoring functions.

**Table S1: Performances on 13 cases of the CAPRI score set.** The number of near-native models among Top10, Top25 and Top200 are reported, showing the good performance of our trained 3D CNN in this task. On most cases the three methods perform similarly with only a few cases where the values differ significantly between the three approaches. iScore seems to perform slightly better than HADDOCK and DeepRank. Note that the models are not clustered, thus we evaluate a high range up to top 200 models. The number in parenthesis in the model column indicates the number of near native conformations.

| | | HADDOCK | | | iScore | | | DeepRank | | |
|---|---|---|---|---|---|---|---|---|---|---|
| | #models | Top10 | Top25 | Top200 | Top10 | Top25 | Top200 | Top10 | Top25 | Top200 |
| **T29** | 1979 (166) | 0 | 0 | **71** | 0 | 0 | 49 | 0 | **2** | 37 |
| **T30** | 1148 (2) | 0 | 0 | 1 | 0 | 0 | 1 | 0 | 0 | 1 |
| **T32** | 599 (15) | 0 | 0 | 0 | **7** | **8** | **12** | 0 | 0 | 2 |
| **T35** | 497 (3) | 0 | 0 | 0 | 0 | 0 | 0 | 0 | 0 | **1** |
| **T37** | 1364 (97) | 0 | 1 | 13 | **4** | **7** | 20 | 2 | 3 | **25** |
| **T39** | 1295 (4) | 0 | 0 | 0 | 0 | 0 | 0 | 0 | 0 | 0 |
| **T40** | 1987 (535) | **10** | **24** | **155** | 0 | 1 | 138 | 1 | 5 | 90 |
| **T41** | 1101 (347) | 1 | 7 | 105 | **10** | **25** | **187** | 5 | 18 | 155 |
| **T46** | 1570 (24) | 3 | 8 | 20 | **6** | **9** | **22** | 1 | 4 | 21 |
| **T47** | 1015 (608) | 10 | **25** | **197** | 10 | 24 | 194 | 9 | 24 | 195 |
| **T50** | 1447 (133) | 1 | **6** | 23 | **2** | 2 | **28** | 0 | 2 | 24 |
| **T53** | 1360 (122) | 5 | **10** | **39** | **6** | 9 | 35 | 1 | 5 | **45** |
| **T54** | 1304 (19) | 0 | 0 | 3 | 0 | 0 | **5** | 0 | 0 | 2 |

#### 3-E Computational Efficiency On Data Generation

As mentioned in the main text, the data generation process is rather efficient within DeepRank. As an example, we compare in **Table S2** the time required to process PDB files in DeepRank and in MaSIF. As seen in this table DeepRank only requires a few seconds per complex while MASIF can go up to about 20 minutes. The experiments are done on Intel(R) Xeon(R) CPU E5-2650 v4 with 125G memory. One CPU was used.

**Table S2. Benchmark of data preprocessing time with DeepRank and MASIF on 123 protein complexes from BM5.**

| **Data Preprocessing Speed** [second/complex] | | |
|---|---|---|
| | **DeepRank** | **MASIF** |
| min | 1.6 | 65.0 |
| max | 14.9 | 1197.0 |
| median | 5.0 | 441.0 |
| mean | 5.7 | 466.7 |
| std | 2.6 | 211.3 |

#### 4- References

1. Lensink, M. F. & Wodak, S. J. Score_set: A CAPRI benchmark for scoring protein complexes. *Proteins Struct. Funct. Bioinforma.* **82**, 3163–3169 (2014).
2. Renaud, N. *et al.* iScore: An MPI supported software for ranking protein–protein docking models based on a random walk graph kernel and support vector machines. *SoftwareX* **11**, 100462 (2020).
3. Geng, C. *et al.* iScore: a novel graph kernel-based function for scoring protein–protein docking models. *Bioinformatics* **36**, 112–121 (2020).
